# Becoming mega-diverse: integrative phylogenomics reveals heterogeneous diversification at high rate in *Astragalus* (Fabaceae)

**DOI:** 10.64898/2026.11.10.765594

**Authors:** Daniele Buono, Diego F. Morales-Briones, Aaron Liston, Shahin Zarre, Gudrun Kadereit

**Affiliations:** Prinzessin Therese von Bayern-Lehrstuhl für Systematik, Biodiversität und Evolution der Pflanzen, Ludwig-Maximilians-Universität München, Munich, Germany; Department of Botany and Plant Pathology, Oregon State University, Corvallis, OR, USA; Department of Plant Sciences, School of Biology, College of Science, University of Tehran, Tehran, Iran; Staatliche Naturwissenschaftliche Sammlungen Bayerns (SNSB): Botanische Staatssammlung München, Botanischer Garten München-Nymphenburg, Munich, Germany

**Keywords:** Leguminosae, Papilionoideae, Angiosperm megadiversity, Diversification rates, Whole-genome duplication, Chromosome evolution, Incomplete lineage sorting

## Abstract

**Background:** Understanding why species diversity is so unevenly distributed across the Tree of Life remains a central question in evolutionary biology. With more than 3,100 species, *Astragalus* (Fabaceae) is the largest genus of flowering plants and provides an exceptional system for investigating the factors of angiosperm megadiversity. However, despite recent phylogenomic advances, uncertainties in phylogenetic relationships and limited integration of available genomic resources have hindered comprehensive evolutionary analyses.

**Methods:** Here, we reconstruct the most comprehensive phylogenomic framework for *Astragalus* to date by integrating newly generated target enrichment data for 127 samples, representing 123 *Astragalus* taxa, with previously published target-capture, transcriptomic, and genome-skimming datasets. The resulting phylogeny includes 407 accessions representing 379 *Astragalus* species and provides a robust backbone for investigating diversification and genome evolution across the genus. Using a representative reduced dataset, we estimated divergence times, quantified diversification-rate heterogeneity, examined phylogenomic conflict, and evaluated evidence for whole-genome duplication (WGD).

**Key Results:** Our analyses recover a largely stable backbone for the deeper Astragalus lineages while revealing extensive gene-tree discordance concentrated in younger radiations, consistent with incomplete lineage sorting and reticulate evolution during rapid Miocene diversification. Diversification analyses indicate sustained lineage accumulation over approximately 18 million years, with pronounced rate shifts in several clades, particularly within the species-rich Neo-Astragalus lineage. Comparisons of alternative WGD scenarios across the phylogeny provided strongest support for a WGD at the crown node of Neo-Astragalus, with alternative placements receiving weaker support. This result provides a testable hypothesis for a WGD near the origin of the most rapidly diversifying lineage in the genus, although chromosome-scale genomic data will be required to confirm its occurrence and timing.

**Conclusions:** Together, our results indicate that the extraordinary diversity of *Astragalus* did not arise from a single evolutionary innovation or explosive radiation. Instead, megadiversity arose through the long-term interplay of sustained diversification, reticulate evolution, genome restructuring, and likely environmental change coupled with ecological adaptation. Beyond providing the most comprehensive phylogenomic hypothesis for the genus, this study establishes a robust framework for future investigations of biogeography, trait evolution, genome evolution, and the processes underlying exceptional species richness in flowering plants.

## Introduction

Species diversity is unevenly distributed across the tree of life and across taxonomic rank. The reasons for this lie in the dynamics of evolutionary and (paleo)ecological change, the unpredictability of phylogenetic constraints and impact of adaptive novelties as well as in the inconsistency and subjective element in species concepts and taxonomy (Benton and Emerson, 2007; De Queiroz, 2007). The genera of flowering plants are no exception to this overall pattern. Although only 28 genera, distributed across 18 families, currently exceed 1000 accepted species, they collectively comprise approximately 13.6% of all flowering plant species (Moonlight et al., 2024). At the other end of the scale stand the 75% of the flowering plant genera that comprise less than five species. Comparative studies suggest that megadiverse angiosperm genera (with more than 1000 species) generally arise from a combination of temporal persistence, ecological opportunities and episodic shifts in diversification rather than through a single explosive radiation (Magallón and Sanderson, 2001; Escudero et al., 2012; Moonlight et al., 2024). Most megadiverse genera are now regarded as monophyletic radiations, not as taxonomic artifacts. Their exceptional diversity has been associated with combinations of long evolutionary persistence, repeated geographic expansion, climatic change, hybridization, and genomic innovation – although the relative importance of these factors varying among clades (Herre et al., 2008; Moonlight et al., 2015; Sigwart et al., 2018; Knapp et al., 2019; Moonlight et al., 2024). However, there is inconsistency of what taxonomists recognize as genera which renders the assignment “mega-diverse genus” somewhat artificial. Probably not all of the potentially megadiverse genera have been recognized as such, due to lack of adequately sampled phylogenetic analyses and the reluctance to merge seemingly well defined genera for the sake of monophyly (Messerschmid et al., 2020; Michelangeli et al., 2022; Kagame et al., 2025). Conversely, some currently recognized megadiverse genera may be split into smaller genera as more comprehensive phylogenetic studies reveal previously unrecognized evolutionary lineages and provide grounds for revising their circumscriptions (Muñoz-Rodríguez et al., 2023).

*Astragalus* L. (Fabaceae, Papilionoideae) represents one of the most extreme examples of plant diversification, comprising over 3100 species (POWO, 2026; Podlech and Zarre, 2013; Barneby, 1964; Fig. 1i) with ca. 1000 more species than the second largest genus (*Taraxacum* F.H.Wigg.) it is in a league of its own (Moonlight et al., 2024). Since Linnaeus established the genus in 1753, its diversity has been documented through more than 270 years of taxonomic work, with a marked acceleration in species descriptions during the nineteenth and twentieth centuries (Fig. 1i). *Astragalus* spans a wide ecological and geographic range predominantly across arid and semi-arid regions in the Northern Hemisphere, but extending into the Andes of South America (Fig. 1ii). Despite being considered the largest genus in flowering plants since the nineteenth century (Frodin, 2004), the evolutionary mechanisms responsible for its extraordinary species richness remain poorly understood, making *Astragalus* an important system for investigating the origins of megadiversity in plants and in general. *Astragalus* appears to follow a similar general pattern as described for most megadiverse genera, with previous studies indicating that its extraordinary richness reflects the interplay of long evolutionary persistence, heterogeneous diversification dynamics, repeated regional radiations, and ecological expansion across arid and montane environments rather than a single burst of diversification (Sanderson & Wojciechowski, 1996; Scherson et al., 2008; Folk et al., 2024a). Elucidating how these processes have interacted across the evolutionary history of the genus, however, depends on the availability of a robust phylogenetic framework with both comprehensive taxon sampling and genome-scale data.

**Fig. 1.**
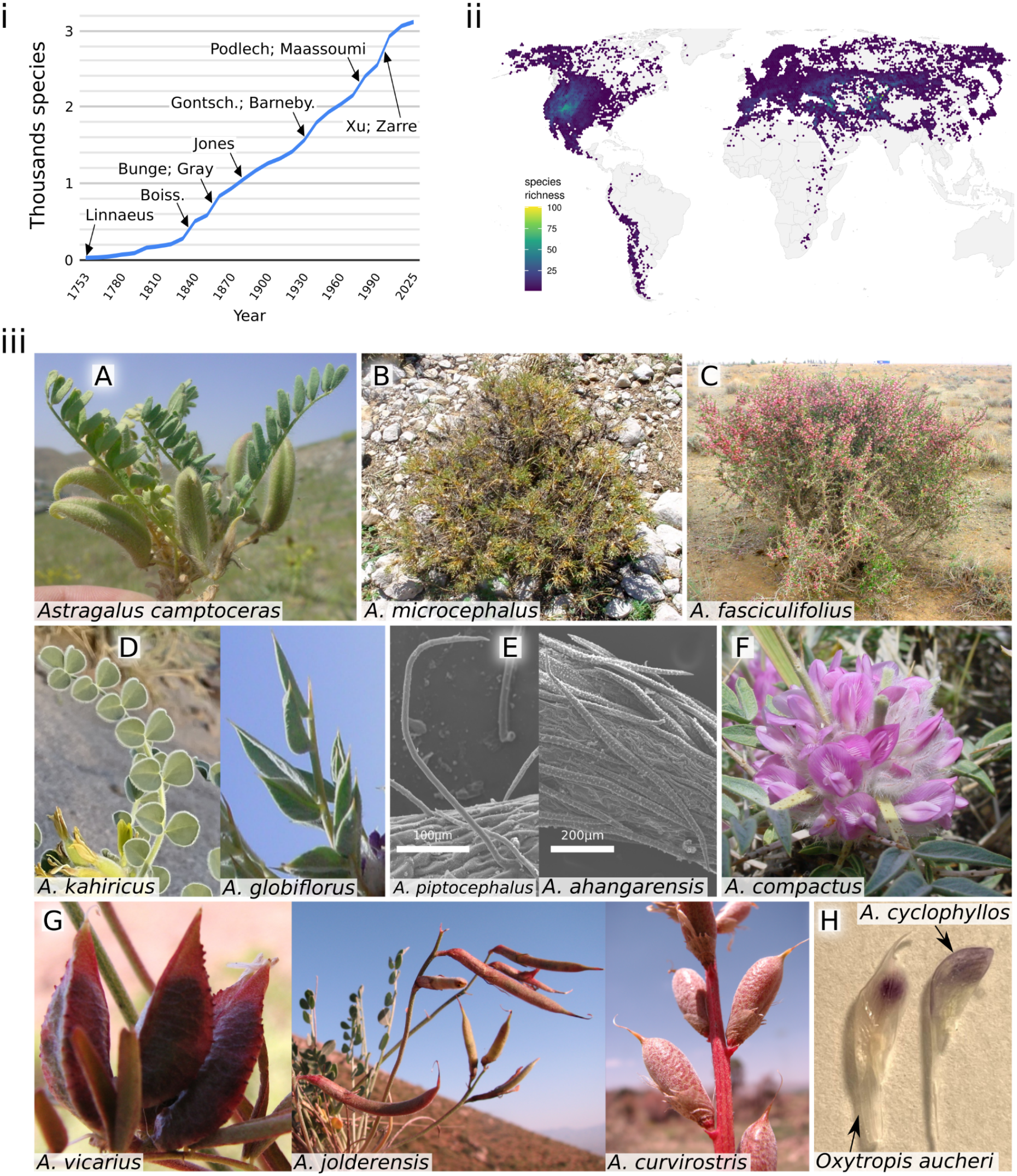
Diversity, distribution, and morphological variation in *Astragalus*. **i)** Increase in the cumulative number of described *Astragalus* species since Linnaeus established the genus in 1753. Arrows indicate the approximate periods of activity of taxonomists who made major contributions to the taxonomy of the genus (not exhaustive), including Linnaeus (1753), Bunge (1864–1884), Barneby (1964), and Podlech and Zarre (2013). Source: World Checklist of Vascular Plants, accessed 23 July 2026. **ii)** Global distribution of *Astragalus*, based on occurrence records, showing its concentration in temperate, arid, and montane regions of the Northern Hemisphere. Species richness is highest in the Irano-Turanian region (>1,500 species), with additional centres of diversity in western North America (∼450 species), the Andes (∼110 species), Central Asia, the Mediterranean Basin, and the Sino-Himalayan region. Source: GBIF, accessed 7 August 2026; additional occurrence data from Munich Herbarium. **iii)** Morphological diversity and diagnostic features of *Astragalus*. **A–C**, growth forms ranging from annual and perennial herbs to cushion-forming plants, subshrubs, and shrubs; **D**, pinnate leaves with entire leaflets; **E**, diverse indumentum, including simple and bifurcate hairs; **F**, papilionoid flowers arranged in axillary racemes; **G**, diverse fruits, commonly dorsally grooved and divided by a complete or incomplete septum; **H**, comparison with the sister genus *Oxytropis*, which is distinguished primarily by its distinctly beaked keel, whereas the keel in *Astragalus* is obtuse or shortly pointed.

Early phylogenetic studies based on single genes or genomic regions established the first broad hypotheses of relationships within *Astragalus* and between the genus and its close relatives (e.g. Sanderson and Wojciechowski, 1996; Wojciechowski et al., 1999; Kazempour Osaloo et al., 2003). These studies provided the initial evidence for the monophyly of *Astragalus* (with a few exceptions), its close relationship with *Oxytropis* DC. and other Old World genera, and the placement of Neo-Astragalus, a New World clade of ca. 500 species, within the broader Eurasian radiation. Neo-Astragalus is characterized by an aneuploid chromosome-number series (n = 11–15), whereas most Old World *Astragalus* species have euploid series based on x = 8, with polyploidy common (Wojciechowski et al., 1999). Building on this foundational work, recent phylogenomic studies have substantially refined our understanding of relationships within *Astragalus*, although important challenges remain. Plastome-based phylogenies generated from genome skimming have clarified relationships among major clades (Su et al., 2021), while transcriptome-scale datasets have provided high-resolution insights into deeper divergences within legumes, including *Astragalus* (e.g., Zhao et al., 2021). More recently, targeted enrichment datasets such as the “NitFix” loci have expanded taxon sampling and enabled broad-scale analyses of diversification and biogeographic history (Folk et al., 2024a), whereas phylogenomic analyses based on hundreds of nuclear loci recovered a highly supported backbone, identified most major clades within the genus and clarified their phylogenetic relationships (Buono et al., 2025a). Together, these studies indicate that the primary phylogenetic structure of *Astragalus* is becoming increasingly well resolved. Nevertheless, substantial uncertainty persists regarding deeper relationships because available datasets differ markedly in genomic scope, taxon coverage, and phylogenetic signal and have largely been analyzed independently, limiting direct comparisons across studies. For example, Buono et al. (2025a) reported extensive gene tree discordance despite strong support for the species tree, implicating incomplete lineage sorting (ILS) and possible hybridization, processes increasingly recognized as major sources of conflict in legume phylogenies (Zhang et al., 2020; Zhang et al., 2026). In contrast, the denser taxon sampling of Folk et al. (2024a), particularly across New World lineages, represents a major advance toward capturing the full diversity of *Astragalus* and refining its biogeographic history, but backbone relationships received comparatively lower support and showed some incongruence with previous phylogenetic hypotheses (Azani et al., 2017, 2019). These discrepancies likely reflect a combination of ILS, reticulate evolution, and paralogy, highlighting the trade-offs between extensive taxon sampling and genome-scale data. Consequently, although the principal lineages and overall backbone structure of *Astragalus* are now emerging with increasing clarity, establishing a robust and stable phylogenetic framework will likely require integrating dense taxon sampling with genome-scale datasets analyzed within a common comparative framework.

Although resolving phylogenetic relationships is essential, understanding the extraordinary diversity of *Astragalus* ultimately requires identifying the evolutionary processes driving its diversification. Geological and climatic processes, such as the uplift of major mountain systems and the expansion of arid environments, have been implicated in promoting diversification through the creation of adaptive opportunity and geographic isolation (Wen et al., 2014). At the same time, genomic factors such as polyploidization and whole-genome duplications (WGD) across angiosperms have been associated with increased genetic novelty, ecological tolerance, and long-term evolutionary success (Soltis et al., 2009; Ren et al., 2018; Heslop-Harrison et al., 2023). Yet, the relationship between WGD and diversification is complex: some studies support a positive association, others highlight a temporal lag between duplication events and subsequent radiations, or even no consistent effect across lineages (Clark and Donoghue, 2018; Landis et al., 2018). In the Astragalean clade (sensu Sanderson and Liston, 1995), widespread variation in chromosome number and ploidy levels suggests a complex history of genome evolution. Whether genome evolution contributed to the exceptional diversification of *Astragalus*, alone or in combination with environmental change and reticulate evolution, remains unknown.

## Goal of the study and research questions

To address inconsistencies among previous data sets, as well as evidence of hybridization, ILS, and paralogy especially in younger branches, we reconstruct a densely sampled, time-calibrated phylogeny of *Astragalus* by integrating multiple genomic datasets, including target enrichment loci, genome skimming, and transcriptomic data. This combined framework enables us to obtain a more comprehensive and representative phylogeny of the genus and its major subclades, establishing a foundation for testing macroevolutionary hypotheses. We investigate whether patterns of diversification coincide geographically and temporally on the phylogeny with inferred WGD events and chromosome evolution. Specifically, we aim to (i) further resolve phylogenetic relationships among major *Astragalus* clades by integrating multiple genomic datasets, (ii) quantify patterns of diversification across the genus and identify potential rate shifts, and (iii) evaluate alternative WGD scenarios and characterize patterns of chromosomal evolution in relation to the diversification history of the genus. By integrating phylogenetic, temporal, and genomic perspectives, this study provides a unified framework for investigating the evolutionary processes associated with the largest radiation in flowering plant genera.

## Material and methods

### Taxon sampling

We designed a sampling strategy to maximize phylogenetic representation across *Astragalus* by targeting all major clades recovered in the most comprehensive and strongly supported studies available to date (Azani et al., 2017, 2019; Su et al., 2021; Folk et al., 2024a; Buono et al., 2025a). To achieve broad coverage of the genus backbone, we compiled a priority sampling list that included representatives of all currently recognized major lineages, with an emphasis on Old World (OW) diversity. This sampling framework comprised 320 OW species, representing ca. 10% of the genus (Table S2). For part of this taxonomic set, molecular data were already available from previous phylogenomic studies based on multiple sequencing strategies, including target enrichment datasets (e.g., Astragalean819: Buono et al., 2025a; NitFix: Folk et al., 2024a), transcriptomic data (Zhao et al., 2021), and genome skimming approaches (Su et al., 2021). To complement these existing resources and fill remaining phylogenetic gaps, we newly generated sequence data for 127 *Astragalus* taxa (Table S1) following the laboratory and bioinformatic protocols described in Buono et al. (2025a, 2025b). To further improve representation of the Neo-Astragalus clade, we additionally incorporated raw sequencing reads from 89 New World (NW) accessions generated by Folk et al. (2024a), selecting samples with the highest sequence recovery and overall data quality. These raw reads were integrated with our newly generated data and other publicly available sequencing data and processed through the same bioinformatic pipeline, including read filtering, locus assembly, and orthology inference, to ensure consistent data processing across studies. The resulting backbone dataset, based on our priority sampling scheme and the integration of newly generated and previously published data, comprised 407 accessions in total, of which 379 belonged to *Astragalus*, representing 347 species. In addition, we assembled a second, expanded dataset including all newly generated samples together with publicly available *Astragalus* data recovered from previous studies (Table S3). This comprehensive dataset comprised 890 accessions in total, including 794 *Astragalus* samples representing 638 species [602 Old World species (ca. 20% of Old World diversity) and 33 New World species (ca. 7% of New World diversity)], and was used to infer a phylogeny with maximized taxonomic coverage across the genus.

### Library preparation, target enrichment, and sequencing

We isolated genomic DNA from approximately 20 mg of dried herbarium material using the NucleoSpin Plant II kit (Macherey-Nagel, Düren, Germany), following the manufacturer’s instructions with minor adjustments. For cell lysis (step 2a), 600 μL of buffer PL1 was applied and samples were incubated for 1.5 h. We omitted RNase A because the age of the specimens made RNA preservation unlikely. In step 6, an additional wash was included by adding 350 μL of buffer PW2 prior to complete membrane drying. DNA was eluted in 50 μL of buffer PE (5 mM Tris-HCl, pH 8.5). Quantification was carried out with the Qubit 4 Fluorometer (Invitrogen) using the High Sensitivity assay kit (Thermo Fisher Scientific, Waltham, MA, USA), and fragment length distribution was assessed on a 1% agarose gel.

For library construction, we adjusted genomic DNA to 250 ng in a final volume of 55 μL buffer PE and sheared to an average fragment size of ∼350 bp using a Covaris M220 Focused-ultrasonicator (Covaris, Woburn, MA, USA). We evaluated fragment size profiles with the Agilent 4150 TapeStation System employing High Sensitivity D1000 ScreenTape (Agilent Technologies, Santa Clara, CA, USA). We prepared sequencing libraries using the NEBNext Ultra II DNA Library Prep Kit for Illumina together with the NEBNext Multiplex Oligos for Illumina (Dual Index Primers Set 1; New England Biolabs, Ipswich, MA, USA), according to the supplier’s recommendations. During adaptor ligation (protocol step 3), we optimized size selection parameters individually based on TapeStation results. We amplified libraries with eight PCR cycles. We quantified final libraries and verified their fragment distributions using the same Qubit and TapeStation procedures described above.

We combined libraries into pools of 25-26 samples, grouping samples with comparable mean fragment sizes and contributing equal DNA amounts (200–250 ng per sample). We vacuum-dried each pool resuspended in 7 μL nuclease-free water prior to target capture. Hybridization-based enrichment was conducted using the myBaits Hybridization Capture Kits (Daicel Arbor Biosciences, Ann Arbor, MI, USA) following protocol version 5.02. We performed both hybridization and wash steps at 60°C. Post-capture amplification consisted of 10 PCR cycles. We purified enriched pools with the NucleoSpin Gel and PCR Clean-up Kit (Macherey-Nagel), and subsequently quantified and assessed for fragment size as outlined above.

Finally, we combined the sample capture pools in equimolar ratios and sequenced by Novogene (Munich, Germany) on an Illumina NextSeq 1000 platform.

### Phylogenomic analysis

CAPTUS v1.5.0 pipeline (Ortiz et al., 2026) was used for data assembly. The cleaning step (captus clean command) was performed using ––trimq 20 to trim leading and trailing regions with PHRED quality score less than 20, and ––maq 20 to keep only reads with minimum average quality of 20. The assembly step (captus assemble) was performed with different settings for the various datasets: ––min_count 10 (minimum contig depth) for NitFix dataset; ––min_count 40 for our data and Buono et al. (2025). For Su et al. (2021) data, the default option for genome skimming data (––preset CAPSKIM) was used. To assemble all sequences targeted by both Astragalean819 (Buono et al., 2025a) and NitFix (Kates et al., 2024), a target file including all sequences targeted by the two bait sets was produced by matching respective *Arabidopsis thaliana* (L.) Heynh. genes, resulting in an integrated target file with 832 loci (819 loci from Astragalean819 plus 13 non overlapping loci from NitFix). The extraction step (captus extract) was performed using the combined target and separately for Astragalaen clade ingroups or outgroups, by setting the minimum percentage of identity to the reference differently for ingroups (nuc_min_identity = 88) and outgroups (nuc_min_identity = 75). We also used a less stringent minimum percentage of coverage of the reference for NitFix (nuc_min_coverage = 10) compared to the other datasets (nuc_min_coverage = 20) after inspecting the number of loci retrieved.

Sequences obtained with CAPTUS were aligned with MACSE v2.07 (Ranwez et al., 2018). Poorly represented alignment columns were filtered using Pxclsq v1.3 (Brown et al., 2017), applying a minimum column occupancy threshold of 10% to exclude sites dominated by missing data. Gene trees were reconstructed with IQ-TREE v3.0.1 (Minh et al., 2020). Substitution models were selected automatically with ModelFinder (option –m TEST; Kalyaanamoorthy et al., 2017), and branch support was assessed with 1,000 ultrafast bootstrap replicates.

To reduce redundancy, tips forming mono-or paraphyletic groups were masked, retaining in each case the sequence with the highest number of unambiguous characters in the trimmed alignment. Outlier branches were pruned using TreeShrink v1.3.9 (Mai and Mirarab, 2018), with a quantile cutoff of 0.1 and excluding designated outgroups from the analysis. Homologous sequence FASTA files were subsequently extracted from the curated trees and realigned with OMM_MACSE v12.01. Homolog gene trees were then inferred again using IQ-TREE under the same settings described above.

Orthology assessment followed the monophyletic outgroup (MO) approach of Yang and Smith (2014). After alignment and cleaning as outlined previously, final ortholog datasets were used to reconstruct gene trees in IQ-TREE with identical model selection and bootstrap parameters. A quartet-based species tree was estimated with ASTRAL v1.23.4.6 (Zhang et al., 2020). In parallel, ASTRAL-Pro3 v1.23.3.6 was employed to infer a species tree from homolog trees, allowing the inclusion of multicopy loci. We used Phytop (Shang et al., 2025) for summarizing the support information provided by ASTRAL analysis. For the concatenation approach, all ortholog alignments were merged into a single supermatrix, which served as input for Maximum Likelihood (ML) species tree estimation in IQ-TREE using the same model testing and bootstrap procedures. After examining the phylogenies obtained, we excluded several samples that were evidently misplaced based on current knowledge or non-monophyly of taxa (flagged in suppl. Table S3).

Patterns of gene tree discordance were examined with PhyParts v0.0.1 (Smith et al., 2015). Prior to this analysis, ortholog trees were rooted with *Medicago truncatula* Gaertn., *Cicer arietinum* L., *Trifolium pratense* L. as outgroups. PhyParts was run twice: first with parameters –a = 0 and –s = 70 to quantify concordant and discordant nodes exceeding a 70% support threshold, and second with –a = 0 to recover nodes below this threshold. Outputs from both runs were combined and visualized in R to calculate the proportions of concordant, discordant, uninformative, and missing data signals.

To reduce missing data and obtain higher support along the backbone of *Astragalus*, the dataset was filtered to keep only samples obtained from Astragalean819 and transcriptome data plus three genomes as outgroup, dropping samples from same taxa, resulting in a dataset of 786 orthologous sequences belonging to 226 taxa including 203 *Astragalus* species. The following analysis is based on this reduced dataset.

### Time-Calibration and Diversification analysis

We performed dating of the ML phylogeny with RelTime, and since no fossil record has so far been confidentially assigned to *Astragalus*, we used two secondary calibration points from the chronogram of Zhao et al. (2021): a root constraint (uniform distribution; minimum: 38.84 million years ago [Mya], maximum: 40.21 Mya), and the split between the Coluteoid clade and the rest of Astragalean clade (uniform distribution; min: 20.63 Mya, max: 21.19 Mya). To investigate shifts in diversification across *Astragalus*, we used BAMM v2.5.0. To adjust for imbalanced sampling across *Astragalus* clades, we estimated our sampling fractions by comparing the phylogenies published by Azani et al. (2017, 2019), Folk et al. (2024a), Buono et al. (2025a); and a large ITS based phylogeny comprising ∼900 *Astragalus* taxa (Azani et al., unpublished results) and known morphological characteristics of the clades. Since the taxonomy of NW uses a broader species concept (Barneby, 1964) compared to the OW (see Discussion), to be able to have the same comparison level between Neo-Astragalus and OW taxa, we also included varieties as distinct taxa in NW (estimated at 726). We also compared the results adjusted for sampling by *Astragalus* clades with an analysis run without adjustment. The input tree included only Astragalean taxa. Recommended priors were produced with the function setBAMMpriors in the package BAMMtools (lambdaInitPrior = 0.48841; lambdaShiftPrior = 0.06372; muInitPrior = 0.48841).

### Gene duplication analysis

To infer gene family counts for internal nodes along the phylogeny, we used Notung v2.9 (Stolzer et al., 2012). We excluded putative recent allopolyploids by removing outliers for number of paralogs in the dataset with the 1.5 * IQR rule using an R script. We also filtered for homolog trees that contained all the taxa (no missing data). We then rearranged homologous gene trees so that poorly supported branches (less than 80% support) were rearranged as in the species tree. To obtain gene counts, we performed reconciliation with the duplication cost set from 1.5 to 5 and the loss cost set to 1.0, thereby reducing the influence of missing data on reconciliation scores. The results of gene counts for tip and nodes and percentages of duplication were then mapped on the species tree using a R script. As the estimation of gene counts across the nodes in the phylogeny were inflated (see below), we mapped instead the number of homolog copies in each clade with the phylogeny, and calculated the likelihood support for different scenarios with WGD happening on the nodes for clades where high proportions of gene copies were observed with the R package WGDgc (Rabier et al., 2014). Birth-death rates and duplicate gene retention were estimated under maximum likelihood using the function MLEGeneCount (Koenen et al., 2021). To be able to use the option oneInBothClades, we removed genes that did not have at least one copy in both main clades at the root. To evaluate alternative whole-genome duplication (WGD) hypotheses, maximum-likelihood gene count models were fitted for each candidate node and compared with a null model lacking a WGD event (Rabier et al., 2014).

Chromosome numbers for *Astragalus* species were gathered from the Chromosome Counts Database (CCDB, https://ccdb.tau.ac.il/, accessed 5 March 2026), the Index to Plant Chromosome Numbers (IPCN, https://legacy.tropicos.org/Project/IPCN, accessed 6 March 2026), *Astragalus* of the world online (http://astragalusofworld.com, accessed 4 March 2026), and Podlech and Zarre (2013), yielding data for 740 *Astragalus* taxa.

## Results

### Sequencing and assembling

We newly generated target enrichment data for 127 samples representing 123 *Astragalus* taxa (Table 1). The number of raw paired-end reads per sample ranged between 1.2 million and 13.4 million (Table S4). Across all samples from all sequencing data sources combined, a total of 738 orthologous sequences were recovered, with a mean of 184.3 orthologs per sample (Fig. S1). Due to poor quality or evident phylogenetic misplacement, some samples were filtered out, resulting in a final dataset of 407 samples (including 379 samples of *Astragalus*). Ortholog recovery varied among data sources. The genome dataset used as outgroups (n = 3) shows the highest recovery by a wide margin, with a mean of 651 orthologous sequences per sample and relatively tight distribution. Transcriptomic data (n = 24; Zhao et al., 2021) yielded an average of 315.9 orthologs per sample. Target enrichment data based on Astragalean819 (Buono et al., 2025a; n = 80) and this study (n = 127), showed a slightly lower but still substantial mean of 273.7 and 251.8 orthologs, respectively, with moderate dispersion. The NitFix dataset (Folk et al., 2024a) (n = 89) showed the lowest recovery, with a mean of 37.3 orthologs per sample and minimal variation. Finally, genome skimming data from Su et al. (2021) (n = 84) exhibited also a lower mean (98.9 orthologs per sample) and a wide dispersion. When all available data were combined into an expanded dataset of 890 accessions across 831 loci (Fig. S3), the number of homologous loci per sample ranged from 1 to 809, and overall matrix completeness was 30.8%.

## Phylogenetic inference

The phylogenetic analyses integrating all filtered datasets recovered largely congruent topologies between concatenation (IQ-TREE) and coalescent-based approaches (ASTRAL and ASTRAL-Pro), particularly along the backbone of the Astragalean clade (Fig. 2). Overall support was high in the concatenated analysis (mean bootstrap, BS = 93.8), whereas coalescent trees showed moderate to high local posterior probabilities (mean LPP = 0.76 and 0.73), with discrepancies primarily concentrated in rapidly diversifying internal clades.

**Fig. 2.**
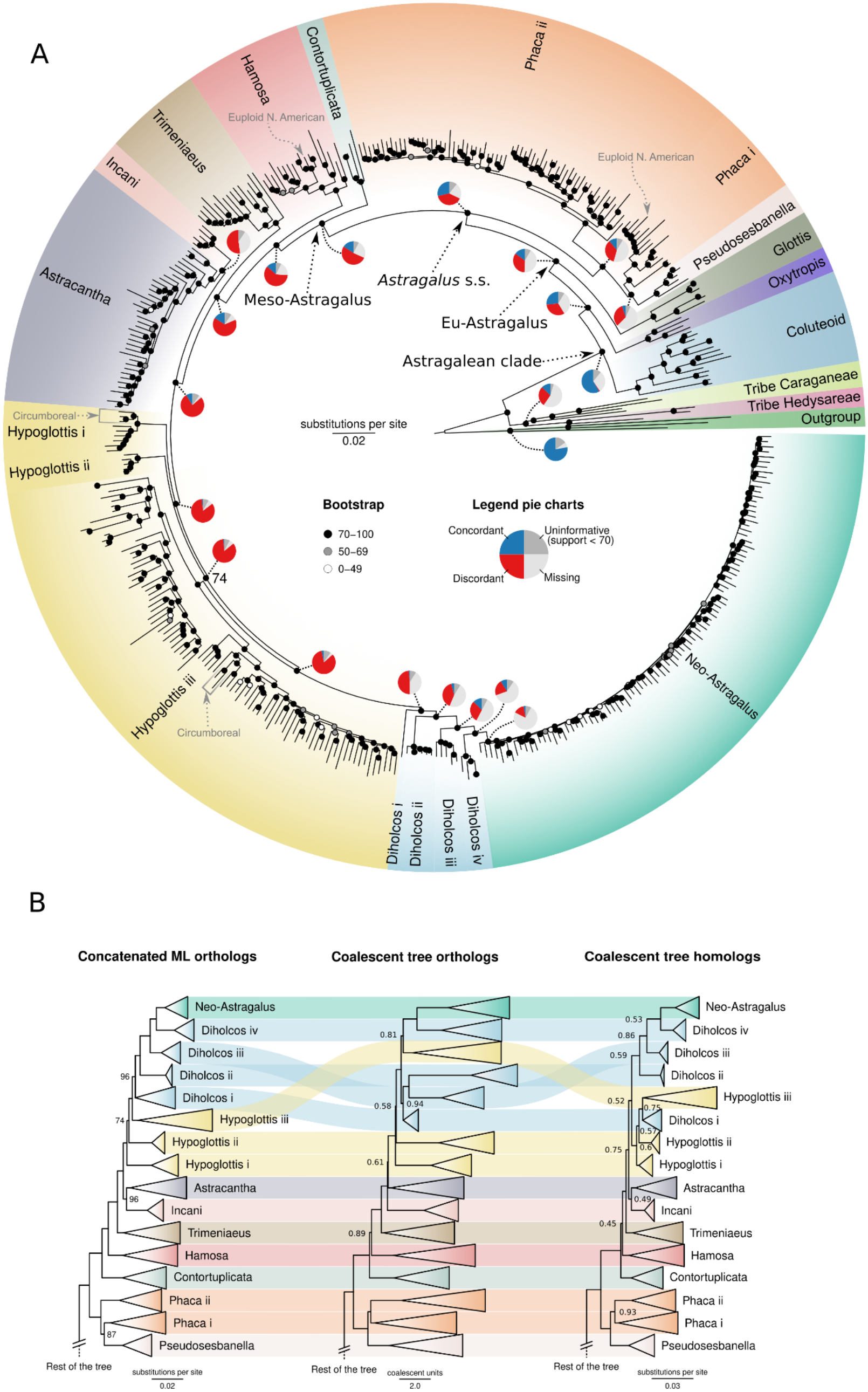
Phylogenomic relationships and comparison of alternative inference approaches across *Astragalus*. The phylogeny includes 407 samples, representing 379 *Astragalus* taxa. **(A)** Circular maximum-likelihood phylogeny inferred from the concatenated dataset, with major recovered clades indicated by different colours. Pie charts along the backbone show gene-tree concordance and discordance inferred using *PhyParts*. Circumboreal and euploid North American species are indicated by grey arrows. Bootstrap support values along the backbone are indicated only when < 95. **(B)** Comparison of topologies inferred using the different methods indicated above each tree. Only the variable portion of the phylogeny, concentrated around the Hypoglottis–Diholcos complex, is shown. Support values are indicated only when Bootstrap < 100 for ML tree and LPP < 0.99 for coalescent trees.

At the deepest nodes, the Astragalean clade, Eu-Astragalus, and *Astragalus* s.s. were consistently recovered as monophyletic with maximal support (BS = 100; LPP = 1). Backbone nodes exhibited low gene tree conflict, as indicated by discordance pie charts dominated by concordant signals (e.g., 430/439 genes supporting the Astragalean clade).

Within *Astragalus*, several major lineages were robustly resolved and monophyletic with maximal support in the ML tree, including Glottis, Pseudosesbanella, Contortuplicata, Hamosa, Trimeniaeus, Incani, Astracantha, and Neo-Astragalus (BS = 100 across nodes; Fig. 2). In contrast, Phaca, Hypoglottis, and Diholcos were consistently recovered as non-monophyletic, with strong bootstrap support for their paraphyletic arrangement in the concatenated tree but markedly reduced support in coalescent analyses (LPP ≈ 0.45–0.81). The node separating Hypoglottis ii and Hypoglottis iii is a notable exception, showing reduced support even in the ML framework.

Patterns of gene tree discordance increase substantially toward shallower nodes, particularly within Meso-Astragalus (sensu Azani et al., 2017), where multiple internal relationships are supported by high bootstrap values despite conflicting underlying gene histories. This is reflected in pie charts showing substantial proportions of discordant and uninformative gene trees at these nodes. Similarly, the clade uniting Eu-Astragalus and *Oxytropis* exhibits elevated conflict (e.g., only 196/433 concordant genes).

The phyTop-based analysis revealed generally low phylogenomic conflict across deep nodes of the Astragalean clade, with low ILS and hybridization index values supporting a largely coherent backbone, including the split between the Coluteoid lineages plus *Oxytropis*, and Eu-Astragalus (Fig. S4). Within *Astragalus*, elevated conflict, attributable to both ILS and hybridization, was concentrated in specific internal regions, particularly around the Hypoglottis and Diholcos complexes.

The ASTRAL-Pro coalescent-based phylogeny, reconstructed from all available *Astragalus* 831 homologous sequences (Fig. S3) following the removal of non-homologous sequences and evidently misplaced samples, comprising 850 tips (760 *Astragalus* samples corresponding to 621 species, of which 588 from Old World and 33 from New World) yielded a backbone topology congruent with that of our filtered dataset (Fig. 2). However, it showed an overall low support (average LPP = 0.579). For this reason, the analyses presented here are based on the filtered dataset.

### Dating and diversification analysis

The dated phylogenies obtained with the combined dataset and high coverage sample dataset showed similar results in terms of clade ages, despite slight differences in the topology recovered (Fig. 3; Fig. S2). The divergence of *Oxytropis* from *Astragalus* was dated at 18.3 Ma (95% CI: 17.81–18.82 Ma), with an age of Eu-Astragalus of 17.93 Ma (95% CI: 17.32–18.57 Ma). *Astragalus* s.s. and Meso-Astragalus originated 15.31 Ma (95% CI: 14.49–16.17 Ma) and 11.73 Ma (95% CI: 10.31–13.37 Ma), respectively.

**Fig. 3.**
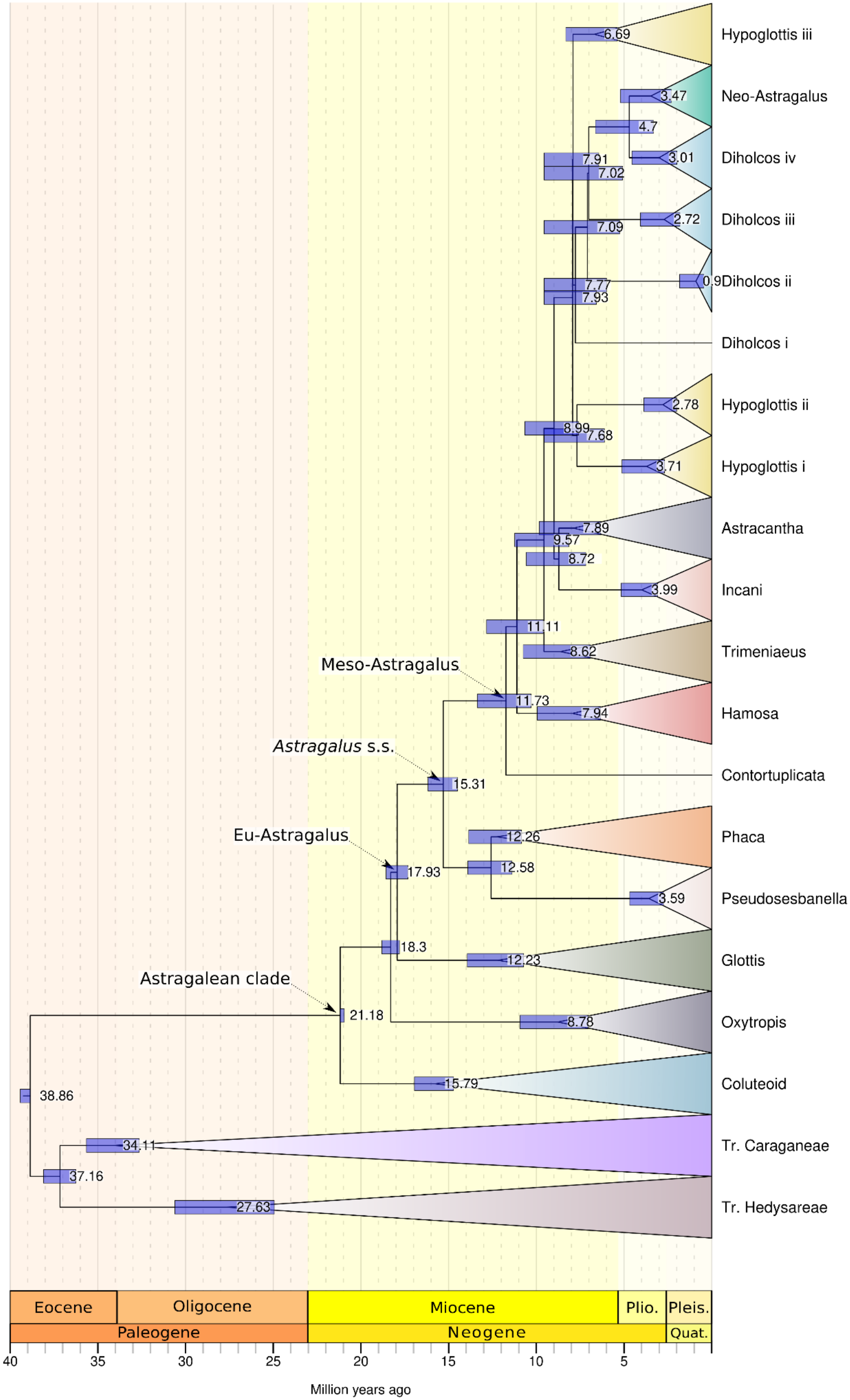
Divergence-time estimation for *Astragalus* and related lineages. The chronogram was inferred using RelTime from a dataset of 786 orthologous loci and 223 taxa, including 203 *Astragalus* species, sampled from Astragalean819 and transcriptomic data. Duplicate samples representing the same taxon were removed. Divergence times were calibrated using secondary age constraints at the root (38.84–40.21 Ma) and at the split between the Coluteoid clade and the remaining Astragalean clade (20.63–21.19 Ma). Blue bars represent 95% confidence intervals for node-age estimates; values indicate median divergence times in millions of years (Ma).

The estimated diversification rates across our dated phylogeny based on BAMM analyses support a scenario of heterogeneous diversification across *Astragalus*, with high rates throughout Eu-Astragalus (mean net diversification across Eu-Astragalus = 0.602) but with clear departures from this background pattern (Fig. 4). The best-supported configuration (f = 0.23; Fig. 4 and S5) identifies three major rate shifts with high posterior probability: (i) within the Phaca clade, with a localized increase in diversification relative to surrounding lineages; (ii) at the divergence of Hamosa from the remaining Meso-Astragalus lineages, indicating an important transition in diversification dynamics, and (iii) a most pronounced shift within the Neo-Astragalus clade, with the highest estimated net diversification rates (approaching ∼2.4), making it the most rapidly diversifying lineage within Eu-Astragalus (Fig. 4). Chromosome number mapped onto the phylogeny reveals additional structure associated with these diversification patterns. In particular, Old World taxa characterized by chromosome numbers of 2n = 16 (euploid) and 2n = 28 (aneuploid), corresponding to the Diholcos iv clade, form a clade sister to Neo-Astragalus.

**Fig. 4:**
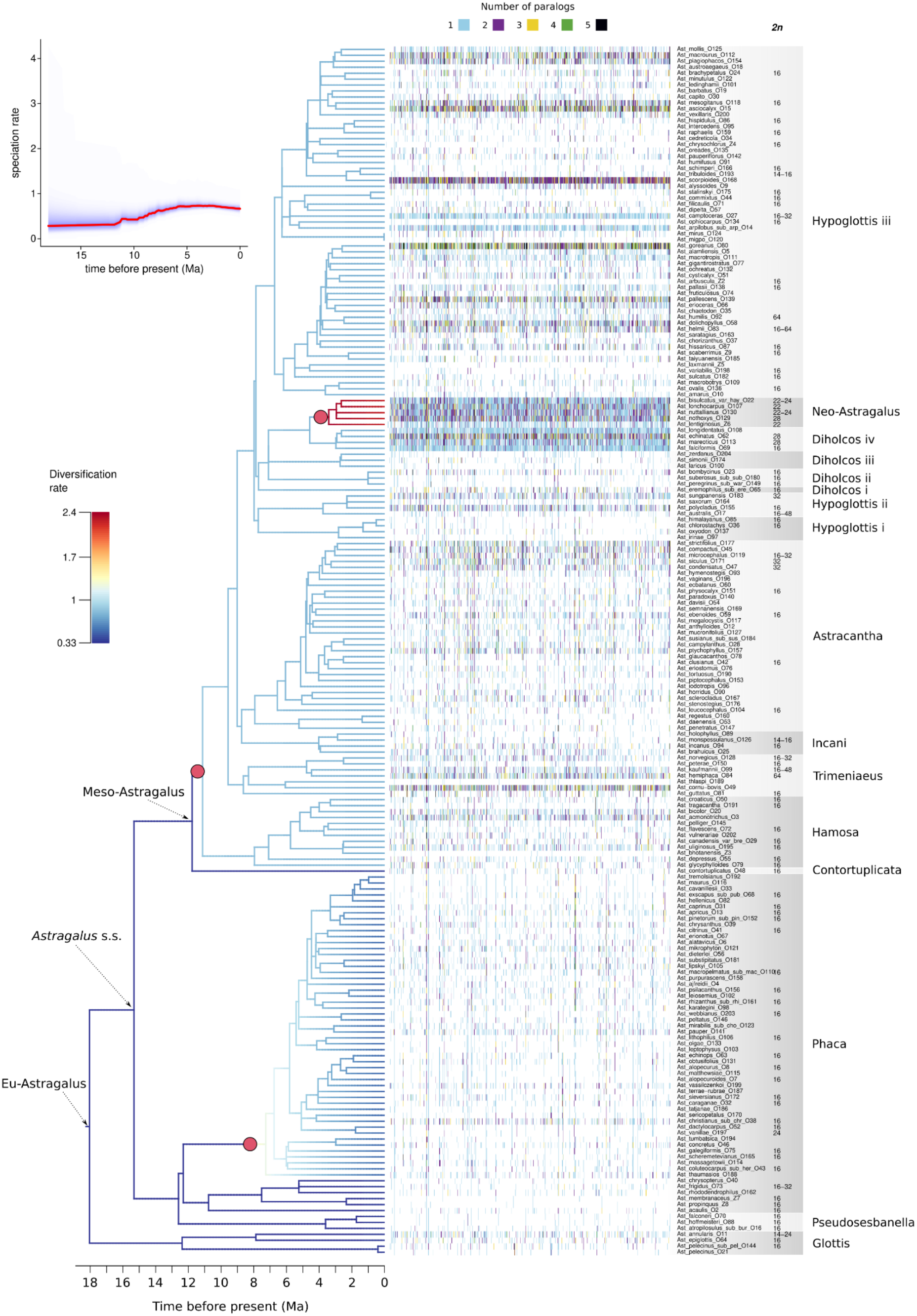
Diversification rates, paralogous gene-copy numbers, and chromosome counts across *Astragalus*. BAMM-inferred rates are shown on the phylogeny, with red circles indicating the three major diversification-rate shifts and the inset on the top left showing the speciation rate following the origin of *Astragalus* 18 Mya. The heatmap shows the number of paralogus gene copies recovered for each sampled taxon, and chromosome numbers (2n), where available, are indicated alongside the phylogeny. The strongest diversification-rate shift occurs within Neo-*Astragalus*, which is also characterized by a distinctive chromosome-number series and elevated numbers of recovered homologous gene copies.

### Exploratory gene-copy and WGD analyses

Gene family reconciliation in Notung (Stolzer et al., 2012), applied here to test the coincidence between genomic restructuring and diversification shifts, inferred extensive duplication events across the phylogeny, with the highest ancestral gene counts reconstructed along the deepest backbone nodes (result not shown). Although duplication signals were detected throughout the tree, ancestral gene counts at several deep nodes were markedly inflated relative to the number of input gene families. Nevertheless, it was still possible to use tip counts to calculate the probability for different WGD scenarios using WGDgc. The model placing a WGD at the Neo-Astragalus crown node provided the best fit among the candidate placements tested (logL = −20,455.29), representing a substantial improvement over the null model (ΔlogL = 191.04; likelihood ratio test (LRT) = 382.08; P < 10^-6^; Fig. S6, Table S5). The estimated duplicate retention rate for this event was 0.406. A WGD placed at the Diholcos iv + Neo-Astragalus node also significantly improved model fit (ΔlogL = 140.57; LRT = 281.14; retention = 0.587), although its likelihood was lower than that of the Neo-Astragalus model. Likewise, a WGD inside the Hypoglottis iii node at the split of *A. camptoceras* was strongly supported relative to the null model (ΔlogL = 66.06; LRT = 132.12; retention = 0.379). In contrast, models placing a WGD at the split between *A. australis* (L.) Lam. with the rest of Hypoglottis ii showed no significant improvement over the null model (ΔlogL = 0.86; LRT = 1.72; P = 0.19), while support for a WGD inside the Astracantha for the node including *A. siculus* Biv., *A. condensatus* Ledeb., *A. microcephalus* Willd., was statistically significant but comparatively weak (ΔlogL = 14.31; LRT = 28.62; retention = 0.055). The model with a WGD inside the Coluteoid clade was indistinguishable from the null model, with an estimated retention rate of zero and identical likelihood values, indicating no evidence for a WGD at this node.

## Discussion

Global analyses have shown that diversification rates are not evenly distributed across angiosperms but tend to be highest in relatively young lineages occupying temperate and seasonally dry environments, whereas the most species-rich tropical floras are largely composed of older, more slowly diversifying clades, although notable exceptions exist (Dimitrov et al., 2023). As a major radiation of the temperate and arid Northern Hemisphere, *Astragalus* is broadly consistent with this pattern. Our analyses suggest that the extraordinary diversity of *Astragalus* has accumulated over approximately 18 million years. With more than 3100 species, *Astragalus* greatly exceeds the diversity of any other flowering plant genus, suggesting that its exceptional species richness is unlikely to be explained solely by the environmental conditions associated with elevated diversification rates at a broad geographic scale.

The key question of why *Astragalus* is so diverse extends to the evolutionary success of the Fabaceae, the third largest family in flowering plants with about 22500 species in 800 genera (Hughes et al., 2025, POWO, 2026). Apart from *Astragalus*, the family includes several large genera, e.g. *Acacia Mill.* (ca. 1084 spp.), *Indigofera* L. (ca. 764 spp.), *Crotalaria* L. (ca. 718 spp.) and *Mimosa* L. (ca. 626 spp.), *Oxytropis* DC. (611 spp.), *Lupinus* L. (585 spp.), and *Inga* Mill. (298 spp.; POWO, 2026). Koenen et al. (2021) dated the crown age of Fabaceae near the Cretaceous-Paleogene (K-Pg) boundary (∼66 mya) and postulated multiple early polyploidization events that coincided with rapid diversification. The fossil record includes a legume fruit assigned to Fabaceae from the upper Campanian of Mexico (∼73.5 mya; Centeno-González et al., 2021), as well as fossil fruits and leaflets from the earliest Paleocene (∼65.35 mya; Lyson et al., 2019). By the middle to late Paleocene, legumes were diverse and abundant in Neotropical rainforest assemblages, with fossils representing several subfamilies (Wing et al., 2009; Herrera et al., 2019). A shift from tropical to temperate regions associated with a shift from long-lived woody growth forms to short-lived mostly herbaceous growth forms resulted in a species-rich lineage, the Hologalegina clade with more than 5000 species, to which *Astragalus* and its sister genus *Oxytropis* belong and constitute c. 75% of its species. Based on this evolutionary history, Hughes et al. (2025) concluded that diversification within Fabaceae has been highly heterogeneous through time and among lineages, with episodic bursts of diversification and turnover rather than a steady accumulation of species. Our results for *Astragalus*, however, reveal a somewhat different pattern. Instead of repeated episodic radiations, we recovered evidence for a sustained diversification over approximately 18 million years at an overall net diversification rate of 0.602. In the context of the global angiosperm radiation, this represents an exceptionally high rate. Magallón and Sanderson (2001) estimated net diversification rates of 0.077–0.089 net speciation events per million years for angiosperms as a whole, based on an estimated standing diversity of 262196 species and an age of approximately 132 my inferred from the fossil pollen record. More recently, Dimitrov et al. (2023) estimated a median genus-level net diversification rate of approximately 0.055 across 14244 angiosperm genera, with the highest values around 0.46. Although methodological differences, including taxonomic scale and estimation approaches, preclude direct quantitative comparisons, our estimate exceeds these rates, placing *Astragalus* among the fastest-diversifying angiosperm genera.

The exceptional diversity of *Astragalus* does not readily point to a single morphological or ecological key innovation. *Astragalus* lacks an obvious trait that could simply explain its unparalleled diversity (Sanderson and Wojciechowski, 1996; Wojciechowski, 2005). A comparable example is provided by *Lupinus*: although the evolution of perenniality has been proposed as a key innovation facilitating montane colonization (Drummond et al., 2012), the repeated radiations of New World *Lupinus* likely reflect the combined effects of ecological opportunities created by mountain uplift, nitrogen-fixing symbioses, and geographic isolation in topographically fragmented landscapes, alongside widespread genome-wide adaptive evolution (Hughes and Eastwood, 2006; Givnish, 2015; Nevado et al., 2016). The exceptional success of *Astragalus* may similarly reflect the cumulative effects of evolutionary innovations inherited from earlier lineages of Fabaceae, including nitrogen-fixing symbiosis and repeated genomic innovation (Faria et al., 2022; Koenen et al., 2020). Within Papilionoideae, traits such as papilionate keel flower, together with ecological versatility and dispersal into new environments in lineages including Hologalegine, may also have contributed (Cai et al., 2025; Hughes et al., 2025). These intrinsic factors may have interacted with ecological opportunities created by Cenozoic cooling, mountain uplift, and the expansion of temperate and arid environments (Zachos et al., 2001). More generally, the importance of putative “key innovations” in driving exceptional diversification has become increasingly debated, as their effects are often difficult to isolate from coincident environmental and historical processes (Rabosky, 2017; Helmstetter et al.; 2023). *Astragalus* exhibit several striking morphological and ecological features, including inflated calyces and legumes (Podlech and Zarre, 2013), deciduous and persistent legumes that facilitate dispersal (Podlech and Zarre, 2013), cushion-forming growth that protects vegetative and reproductive buds from cold (Hardion et al., 2016), production and/or accumulation of alkaloids and nitrotoxins against herbivory (Wink, 2013), edaphic specialization to marginal habitats (Jones et al., 2021), and spines (Hardion et al., 2016). Many of those features have evolved or been retained in different parts of the genus. Whether any of these traits contributed to diversification remains an open question requiring explicit trait-dependent analyses. Thus, rather than identifying a single innovation responsible for its exceptional richness, our results point to a history of sustained diversification within a lineage whose ecological and evolutionary characteristics may have facilitated repeated opportunities for diversification (Donoghue and Sanderson, 2015).

### Rapid radiation, phylogenomic conflict, and Miocene diversification

Although the backbone of *Astragalus* appears relatively stable, our analyses also reveal extensive phylogenomic conflict concentrated in younger, more recent radiations, particularly within Meso-Astragalus and within the Hypoglottis and Diholcos lineages (Fig. 2). This conflict is expressed by the marked discrepancy between the high bootstrap support recovered in concatenated ML analyses and the lower local posterior probabilities observed in coalescent analyses (Fig. 2B). Such patterns are now widely recognized in phylogenomics and often reflect the inability of concatenation methods to adequately accommodate heterogeneous evolutionary histories among loci (Kubatko and Degnan, 2007; Roch and Steel, 2015). Consistent with this interpretation, the discordance pie charts and PhyTop analyses (Fig. 2A, S4) suggest that these unstable regions are characterized by elevated ILS and moderate signals of hybridization, whereas deeper nodes exhibit comparatively little conflict, similarly to what was reported in Buono et al. (2025a).

The concentration of discordance toward shallow nodes is consistent with rapid successive speciation events, during which short internodes may have limited the sorting of ancestral polymorphisms (Maddison, 1997; Degnan and Rosenberg, 2009; Mallet et al., 2016; Morrison, 2016). Similar patterns have been documented repeatedly in recent angiosperm radiations and across papilionoid legumes more broadly (Pease et al., 2016; Koennen et al., 2020; Morales-Briones et al., 2021; Choi et al., 2022; Zhang et al., 2026). In particular, the concentration of discordance around the Diholcos and Hypoglottis complexes may reflect particularly rapid diversification and/or historical introgression events associated with ecological expansion across Eurasian and circum-boreal regions. The relatively weak support recovered even in concatenated analyses for some relationships within these complexes further indicates that the underlying evolutionary history may be difficult to represent with a single bifurcating topology, as also suggested by previous analyses (Buono et al., 2025a).

The temporal framework inferred here places this diversification within a broader period of Neogene environmental change. Divergence-time analyses suggest that the split between *Astragalus* and *Oxytropis* occurred approximately 18.3 Ma, followed shortly by the origin of Eu-*Astragalus* at 17.9 Ma, both during the Early Miocene. The subsequent origin of *Astragalus* s.s. (15.3 Ma) and especially Meso-Astragalus (11.7 Ma) occurred during the Middle Miocene and near the Middle-Late Miocene transition, respectively. These intervals fall within a broader period of climatic, tectonic, and ecological reorganization across Eurasia. More specifically, aridification of the Asian interior was already underway by the Early Miocene (Guo et al., 2002). During the Middle Miocene, uplift of the Tibetan Plateau and adjacent mountain ranges further increased topographic and climatic heterogeneity, while steppe vegetation began to spread in Central Asia around 15 mya (Barbolini et al., 2020). A major global climatic transition also occurred during this period, which was characterized by cooling and expansion of the East Antarctic ice sheet (Flower and Kennett, 1994). Previous studies have similarly linked *Astragalus* diversification to Neogene aridification, biome shifts, and the expansion of open habitats (Azani et al., 2017, 2019; Folk et al., 2024a).

Taken together, our results are consistent with a history of rapid diversification characterized by both ILS and reticulate evolution. The discordance detected here, together with network-based evidence from previous work (Buono et al., 2025a), highlights the need for denser sampling and explicit network analyses to test the extent and evolutionary significance of reticulate histories across *Astragalus*. Such analyses may ultimately provide a complementary perspective to bifurcating species-tree approaches, particularly in the youngest and most rapidly diversifying lineages of the genus (Nakhleh, 2010; Yu and Nakhleh, 2015).

### Diversification-rate heterogeneity and chromosome-number evolution

Within this Miocene environmental context, diversification analyses suggest that *Astragalus* represents a mega-radiation characterized by sustained background diversification combined with localized accelerations rather than a single explosive radiation. Similar patterns have recently been documented across flowering plants and within *Astragalus* itself (Tank et al., 2015; Folk et al., 2024a). The most pronounced shift occurs within the Neo-Astragalus clade, comprising more than 700 taxa (species, subspecies and varieties; Barneby, 1964; Johnston, 1938; Johnston, 1947; Scherson et al., 2008). Here, infraspecific taxa are treated as species-comparable entities for better congruence with the taxonomic concept applied in Old World *Astragalus* (Podlech and Zarre, 2013). Weaker, but still pronounced, accelerations occur near the divergence of Hamosa and within Phaca clades. These results indicate that diversification heterogeneity is distributed across several independent lineages, producing a mosaic pattern of radiation rather than a uniform increase in speciation throughout the genus.

The chromosomal patterns recovered here provide additional insights into the mechanisms that may underlie these lineage-specific radiations. Neo-Astragalus, which contains the vast majority of New World species, is characterized by a derived aneuploid chromosome series with haploid numbers ranging from *n* = 11–15 (Wojciechowski et al., 1999). In contrast, the comparatively few North American species (around 11-13) that fall outside Neo-Astragalus, including several circumboreal taxa and other independently derived lineages, retain the ancestral euploid chromosome condition and are found nested within multiple Eurasian clades (or groups) in the phylogeny. Our phylogenetic results are consistent with the hypothesis that the members of Neo-Astragalus share a common ancestor in which chromosome-number restructuring had already occurred, whereas the euploid American taxa represent several independent colonization events that did not subsequently give rise to major radiations.

This contrast is notable because the exceptional diversification of Neo-Astragalus coincides with both a shared history of chromosome-number evolution and the strongest diversification-rate shift recovered in our analyses. The sister relationship between Neo-Astragalus and the Diholcos iv clade, together with elevated numbers of homologous gene copies inferred in this region of the phylogeny, provides an additional signal of distinctive genomic history in this lineage (Fig. 4). When alternative WGD scenarios were evaluated across the backbone, the scenario placing a WGD prior to the origin of Neo-Astragalus received the highest probability. Taken together, the coincidence of chromosome-number evolution, elevated recovered copy counts, the inferred WGD scenario, and the strongest diversification shift identifies the origin of Neo-Astragalus as a particularly important target for future genome-scale investigation. However, these patterns do not establish a causal relationship between WGD, chromosome-number evolution, and diversification.

Polyploidy and WGD have repeatedly been associated with diversification and ecological expansion in angiosperms and legumes (Tank et al., 2015; Van de Peer et al., 2017; Ren et al., 2018), although their macroevolutionary consequences remain debated, with several studies emphasizing delayed or lineage-dependent effects (Landis et al., 2018; Heslop-Harrison et al., 2023; Hagen and Beaulieu, 2024). In *Astragalus*, the contrasting histories of North American lineages provide an intriguing comparative context: several euploid lineages independently colonized North America without undergoing comparable diversification, whereas the aneuploid Neo-Astragalus lineage subsequently underwent extensive species proliferation. This radiation also extends into South America, where approximately 110 species of *Astragalus* occur, with major centers of diversity in the Andes (Scherson et al., 2008). This pattern suggests that dispersal into the New World alone was insufficient to generate megadiversity and raises the hypothesis that the exceptional diversification of Neo-Astragalus may have resulted from an interaction between ecological opportunity and lineage-specific genomic history, including WGD and subsequent chromosome-number changes. In particular, aneuploidy may have contributed to the ability of Neo-Astragalus to exploit novel environmental opportunities by generating genomic variation potentially facilitating adaptation to different ecological conditions, thereby promoting rapid diversification. However, chromosome-number changes have also occurred sporadically in otherwise predominantly euploid Old World lineages. For example, reduced basic chromosome numbers of x = 7 and x = 6 have been reported in several Old World species, including *A. annularis* Forssk. (2n = 14) and *A. boeticus* L. (2n = 30, pentaploid based on x = 6; Badr et al., 1996; Badr and Sharawy, 2007). These occurrences suggest that chromosome-number changes alone may not be sufficient to promote increased diversification, but that their evolutionary consequences may depend on the ecological context in which they occur and the evolutionary history of the affected lineages. Testing this hypothesis will require chromosome-scale genome assemblies and broader genomic sampling across Neo-Astragalus and its closest relatives.

This interpretation is consistent with the extensive cytogenetic diversity documented across *Astragalus* and *Oxytropis*, where both auto– and allopolyploidization appear common (Záveská et al., 2019; Sandanov et al., 2023). However, our analyses also highlight the methodological challenges of inferring genome evolution from heterogeneous target-enrichment datasets. Duplication–loss reconciliation in Notung (Stolzer et al., 2012) produced extensive duplication signals concentrated along the deep backbone of the phylogeny, yielding ancestral gene counts that greatly exceeded the number of analyzed gene families. Given the pervasive gene tree discordance and strong signatures of ILS recovered across the genus, these reconstructions likely reflect the tendency of reconciliation methods to interpret topological conflict as repeated duplication and loss events rather than genuine large-scale genome duplications, as demonstrated in other systems (Hahn, 2007; Rasmussen and Kellis, 2012). Consequently, the inferred ancestral gene counts should be interpreted cautiously and primarily as indicators of extensive phylogenomic conflict rather than direct estimates of ancestral genome content. Resolving the relationship between chromosome-number evolution, WGD, and diversification will therefore require chromosome-scale genomic resources and analytical approaches that can jointly accommodate gene duplication, ILS, and reticulate evolution.

### *Astragalus* as a model of angiosperm megadiversity

The diversification history inferred here aligns with emerging views on the origin of megadiverse angiosperm genera. Comparative studies indicate that exceptional species richness rarely results from a single evolutionary innovation or a brief episode of explosive radiation, but instead reflects the cumulative effects of environmental change, geographic expansion, niche availability, and lineage-specific genomic processes acting over extended timescales (Magallón and Sanderson, 2001; Escudero et al., 2012, Moonlight et al., 2024).

Comparable multifactorial diversification histories have been documented across numerous megadiverse plant genera. In *Ficus* L., diversification has been linked to long-term coevolution with highly specialized pollinating fig wasps, repeated geographic expansion, and ecological niche partitioning that promoted lineage persistence and regional radiations (Herre et al., 2008). In *Solanum* L., another big genus of flowering plants, diversification appears to have resulted from repeated colonization of novel environments, extensive morphological innovation, and multiple regional radiations across the Neotropics, Australia, and Africa (Knapp et al., 2019). Although *Quercus* L. does not qualify as megadiverse genus, its substantial species diversity (477 spp., POWO, 2026) has been attributed to extensive hybridization and introgression, ecological divergence, and climatic oscillations that facilitated both lineage persistence and repeated episodes of diversification across the Northern Hemisphere (Liu et al., 2025). In *Begonia* L., phylogenetic and biogeographic studies suggest that species richness arose through repeated dispersal events followed by regional radiations, particularly in montane tropical regions, where geographic isolation, habitat heterogeneity, and high rates of local endemism promoted rapid diversification (Moonlight et al., 2015; Moonlight et al., 2024).

Rather than representing an exceptional case, *Astragalus* appears to exemplify a general macroevolutionary pathway by which extreme species richness evolves in flowering plants. Our results suggest that its extraordinary diversity was not generated by a single key innovation or an isolated burst or rapid speciation, but instead emerged through the cumulative interaction of sustained diversification over nearly 18 million years, Miocene environmental change, repeated ecological expansion, reticulate evolution, and lineage-specific genomic restructuring. This combination of processes closely parallels patterns inferred for other megadiverse angiosperm genera, while providing one of the clearest examples in which these mechanisms can be examined within a robust phylogenomic framework. As the largest genus of flowering plants, *Astragalus* therefore represents an important model system for understanding how long-term interactions between environmental opportunity and intrinsic evolutionary processes generate exceptional biodiversity.

### Biogeographic implications

The phylogenetic placement of Neo-Astragalus recovered here is consistent with previous biogeographic hypotheses proposing a colonization of the New World from Old World ancestors (Barneby, 1964; Sanderson and Wojciechowski, 1996; Azani et al., 2017, 2019). Although we did not undertake an explicit historical-biogeographic analysis, the phylogenetic relationships recovered in this study are consistent with a history involving multiple transcontinental exchanges within *Astragalus* rather than a single colonization event. The earliest of these appear to represent exchanges of temperate and boreal taxa between Eurasia and North America, probably via the Bering land bridge during the Miocene, best exemplified by circumboreal species such as *A. laxmannii* Jacq., *A. australis*, and *A. alpinus* L. The position of Neo-Astragalus is also compatible with the previously proposed scenario of a later dispersal from a Mediterranean–West Asian lineage into western North America, followed by extensive diversification (Azani et al., 2019). The mechanism underlying this colonization remains unknown, but zoochory by migratory birds represents one possibility. Seeds of *A. contortuplicatus* L., for example, can survive passage through mallard digestive tracts and subsequently germinate, with potential dispersal distances estimated at up to 1600 km (Lovas-Kiss et al., 2015). This proposed history would differ from the earlier circumboreal migrations because it involves the establishment of a lineage in a novel region followed by rapid diversification, rather than the expansion of already cold-adapted taxa across a continuous high-latitude biome (Donoghue and Sanderson, 2015). The timing of this colonization coincides broadly with the Late Miocene–Pliocene development of increasingly heterogeneous, seasonally dry landscapes in western North America, providing ecological opportunities that may have promoted their subsequent radiation (Zachos et al., 2021; Folk et al., 2024b).

The ecological characteristics of the ancestral lineage also remain uncertain because we did not reconstruct ancestral ecological traits. Nevertheless, the possibility that the ancestor of Neo-Astragalus possessed ecological tolerances suited to seasonal drought, high climatic variability, or open habitats is compatible with recent evidence that diversification and endemism in *Astragalus* are associated with dry montane environments, precipitation seasonality, and isothermality, and that climatic niche characteristics can show phylogenetic conservatism in the genus (Folk et al., 2024a, 2024b). Such preadaptations may help explain why earlier circumboreal dispersals resulted in relatively limited diversification despite repeated exchanges between Eurasia and North America. Under this hypothesis, the exceptional radiation of Neo-Astragalus reflects not dispersal alone, but the interaction between ecological opportunity and a lineage already predisposed to exploit it, consistent with the concept of synnovation (Donoghue and Sanderson, 2015).

The exceptional diversification of Neo-Astragalus may therefore reflect the combined effects of dispersal-mediated ecological opportunity, ecological predisposition to dry and climatically seasonal environments, geographic isolation across western North America, and intrinsic genomic factors, including chromosomal evolution, polyploidization, and reticulation. Available published chromosome counts provide no evidence for additional polyploidization in extant Neo-Astragalus lineages following the proposed ancient event. Our WGDgc analysis provide a specific genomic hypothesis in this regard: among the nodes tested, the best fitting model placed a candidate WGD at the crown node of Neo-Astragalus, whereas alternative placements at the Diholcos iv + Neo-Astragalus node received weaker support and most other tested nodes showed little or no support. This result should therefore be interpreted as a candidate placement of an ancient WGD rather than as evidence that genome restructuring occurred at the Neo-Astragalus crown. Distinguishing an ancient polyploidization event from other sources of gene duplication, and establishing its genomic consequences, will require broader gene-family sampling and independent validation using chromosome-scale genome assemblies, comparative synteny, and related genomic approaches. Thus, the present analysis identifies a testable hypothesis concerning the genomic history of Neo-Astragalus rather than providing definitive evidence for an early episode of genome restructuring.

### Integrative phylogenomics and methodological considerations

Recent studies have demonstrated that target enrichment, transcriptomic, and genome-skimming datasets can be successfully integrated to improve taxonomic breadth and phylogenetic resolution, even when datasets differ substantially in locus composition, sequencing depth, and capture design (Pezzini et al., 2023; Buono et al., 2025a, Chen et al., 2026). Our study follows this emerging trend by integrating data derived from multiple bait sets (Astragalean819 and NitFix), transcriptomic assemblies, and genome-skimming resources into a single large-scale phylogenomic framework for *Astragalus*.

Despite the methodological complexity inherent to merging heterogeneous datasets, the resulting phylogenies recovered a stable deep backbone, with strong support for the Astragalean clade, Eu-Astragalus, and *Astragalus* s.s., consistent with recent phylogenomic reconstructions (Azani et al., 2019; Su et al., 2021; Folk et al., 2024a; Buono et al., 2025a). The high proportion of concordant loci recovered for these deeper nodes gained across different datasets further indicates that ancient divergences within the group represent phylogenetically robust lineages. These lineages retain strong phylogenetic signals despite extensive taxonomic and genomic heterogeneity within those clades.

Nevertheless, several methodological and biological factors remain important sources of uncertainty in large-scale phylogenomic syntheses. Differences in bait design, sequencing strategy, assembly pipelines, missing data, and orthology inference can strongly influence locus recovery and downstream analyses (Vatanparast et al., 2018; Joyce et al., 2025). By including more samples compared to Buono et al. (2025a), and using the same phylogenetic approach to exclude paralogous sequences, the average number of orthologous sequences per each newly sequenced sample was lower (251.8 vs 621.7), highlighting the challenges in inferring paralogs when larger datasets possibly include alloploid individuals. This is also reflected in the lower support along the phylogeny backbone obtained when including all the available sequences (Fig. S3). In addition, merging public datasets introduces uncertainties related to taxonomic misidentification, incomplete voucher documentation, and metadata quality.

These issues are particularly relevant in *Astragalus*, where species delimitation is often difficult due to extensive morphological convergence, narrow endemism, and the existence of diverging taxonomic frameworks for the western and eastern hemispheres (e.g. Barneby, 1964; Podlech and Zarre, 2013). Barneby employed a broad species concept for North America, reducing many previously described species to varieties when morphological differences were minor (e.g. differences in size) or if intermediate plants were documented.

He recognized 368 species and 254 varieties (552 taxa) in North America. Most of these varieties had been previously treated more narrowly and as distinct species (Barneby, 1964). In the most extreme example, this led to nearly 50 recognized varieties of the widespread *A. lentiginosus* Douglas. In contrast, traditional *Astragalus* taxonomy in the Eastern Hemisphere has generally recognized a larger number of narrowly circumscribed species, as exemplified by the treatment in *Flora of the USSR* (Gontscharov and Borissova, 1946). Modern Russian taxonomic literature continues to follow this treatment (Grossheim, 1952; Abdullaeva et al., 1981). Podlech and Zarre (2013) reassessed the numerous taxa described in Soviet literature, and synonymized many names. In this context, many currently recognized species may still represent morphological taxa rather than independently evolving evolutionary lineages. As an example, the recently described *A. dalahuensis* Avazzadeh, Bagheri & Maassoumi, shows clear morphological differences from *A. bruguieri* Boiss. and *A. russellii* Banks & Sol., yet the phylogenetic signals supporting their separation are not significant (Avazzadeh et al., 2025). In contrast, a growing number of population genetic studies have confirmed a higher phylogenetic divergence and the recognition of additional formal species within certain species complexes, despite only minor morphological differences between them (Maylandt et al., 2025; Záveská et al., 2019).

### Synthesis and Future directions

Collectively, our results provide a phylogenomic framework for investigating the evolutionary history and exceptional diversity of *Astragalus*. Rather than demonstrating a single mechanism underlying megadiversity, this study identifies several processes and patterns that warrant further investigation. We document heterogeneous diversification across the genus, extensive phylogenomic discordance consistent with a complex reticulate evolutionary history, and substantial variation in chromosome number based on published cytological data. Our WGDgc analyses further nominate a candidate WGD near the crown of Neo-Astragalus, although this hypothesis requires validation with chromosome-scale genome assemblies, synteny, and broader gene-family sampling. Together, these results provide a basis for testing how environmental change, lineage diversification, reticulate evolution, and genome evolution may have contributed to the exceptional richness of *Astragalus*, but do not by themselves demonstrate that these processes jointly generated its megadiversity.

Despite providing broad taxonomic and genomic coverage, substantial gaps remain, particularly among geographically restricted Old World taxa. Expanding taxon sampling is therefore a major priority; we provide a prioritized list of key taxa whose inclusion is expected to improve phylogenetic resolution and backbone stability (Table S2). Given the enormous size and geographic breadth of the genus, achieving near-complete sampling will require coordinated efforts across collections and research groups.

A major lesson from this study is the continuing importance of herbarium collections as genomic resources. Much of the diversity of *Astragalus* is concentrated in regions where field sampling can be difficult because of political or logistical constraints, making historical collections an essential source of taxonomic diversity. Our extensive use of material from the Botanische Staatssammlung München (M) illustrates the value of natural history collections for expanding phylogenomic sampling. Continued improvements in sequencing technology should make it increasingly feasible to incorporate poorly sampled, endemic, and potentially extinct lineages from historical material.

Our results also highlight the challenges of integrating heterogeneous phylogenomic datasets. Although combining target enrichment, transcriptomic, and genome-skimming data substantially increased taxonomic representation, differences in locus recovery, sequencing strategy, and extensive incomplete lineage sorting limited phylogenetic resolution in several parts of the phylogeny. Future large-scale studies would therefore benefit from standardized sequencing using dedicated probe sets, such as Astragalean819 (Buono et al., 2025b), to generate a dense and well-resolved phylogenomic backbone. This framework will enable more rigorous tests of historical biogeography, diversification, trait evolution, and ecological niche evolution.

Finally, genome evolution remains one of the most promising avenues for future research. Expanded chromosome counts, chromosome-scale assemblies, and population genomic data from representatives of the major clades identified here will be essential for disentangling the contributions of WGD, dysploidy, hybridization, and small-scale duplication to diversification. Integrating these resources with increasingly complete phylogenetic sampling promises not only to clarify the evolutionary history of *Astragalus*, but also to establish the genus as one of the premier model systems for investigating the origins of angiosperm megadiversity.

## Author contributions

D.B., D.M.B., and G.K. conceived the idea. D.B. and D.M.B. performed the experiments and conducted formal analyses. S.Z. and A.L. contributed to species identification and critical revisions. All authors contributed to the discussion and interpretation of the results. D.B. wrote the original draft of the manuscript and all authors contributed to the final draft. G.K. acquired funding and provided resources.

## AI Assistance

ChatGPT was used as an aide in the writing of this manuscript.

## Availability of Data and Materials

Target enrichment data generated for this study can be found in the NCBI BioProject PRJNA1521383 (see Table S1 for SRA accessions). Analyses files are available from the Dryad repository DOI: 10.5061/dryad.ncjsxktc4.

## Acknowledgements

We thank Nasim Azani, University of Tehran (Iran), for sharing the preliminary ITS-based phylogeny of *Astragalus* that was used as reference to assess clade coverage.

## Supporting Information

**Fig S1:**
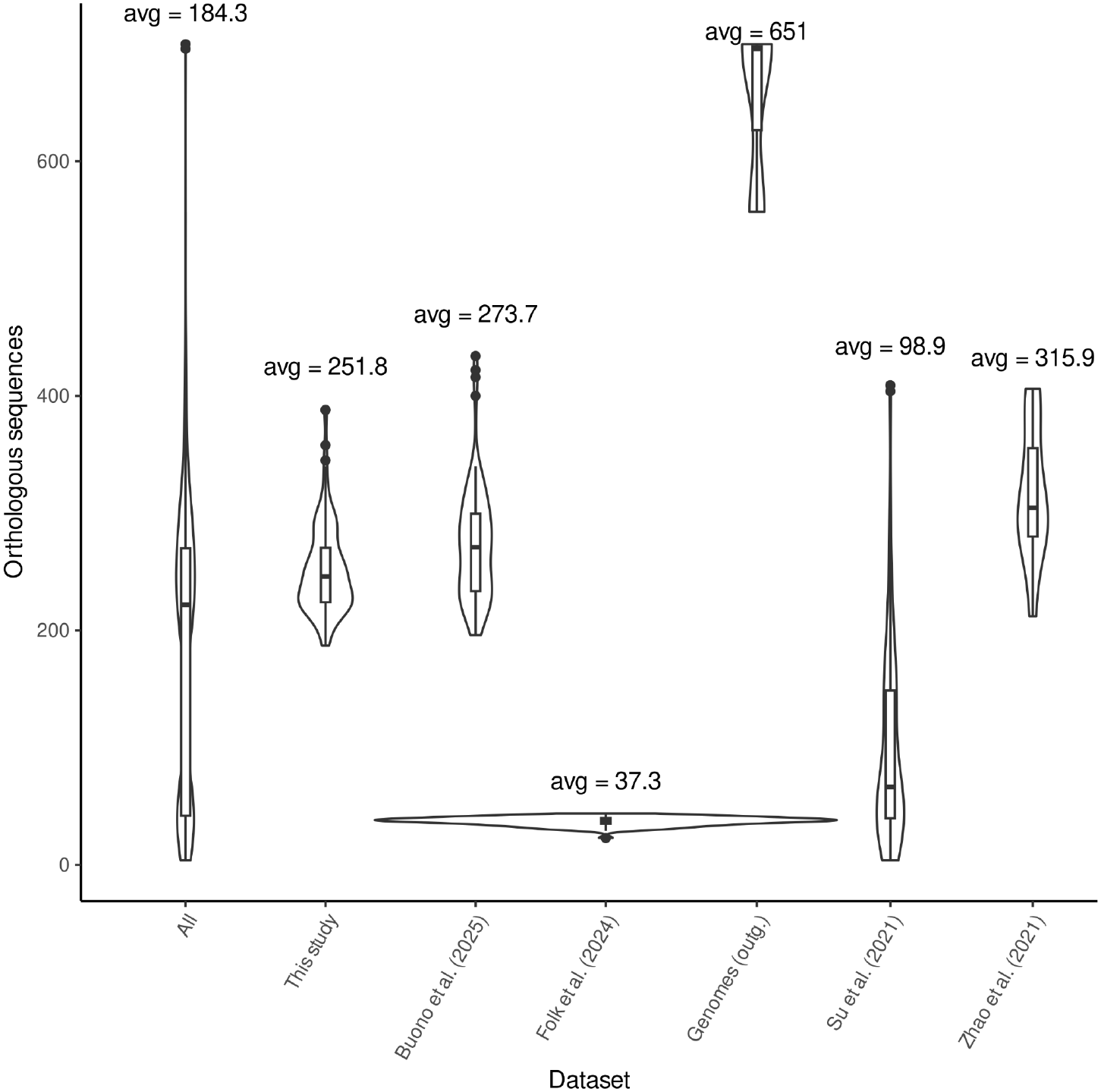
orthologous sequences inferred from different data sources.

**Fig. S2:**
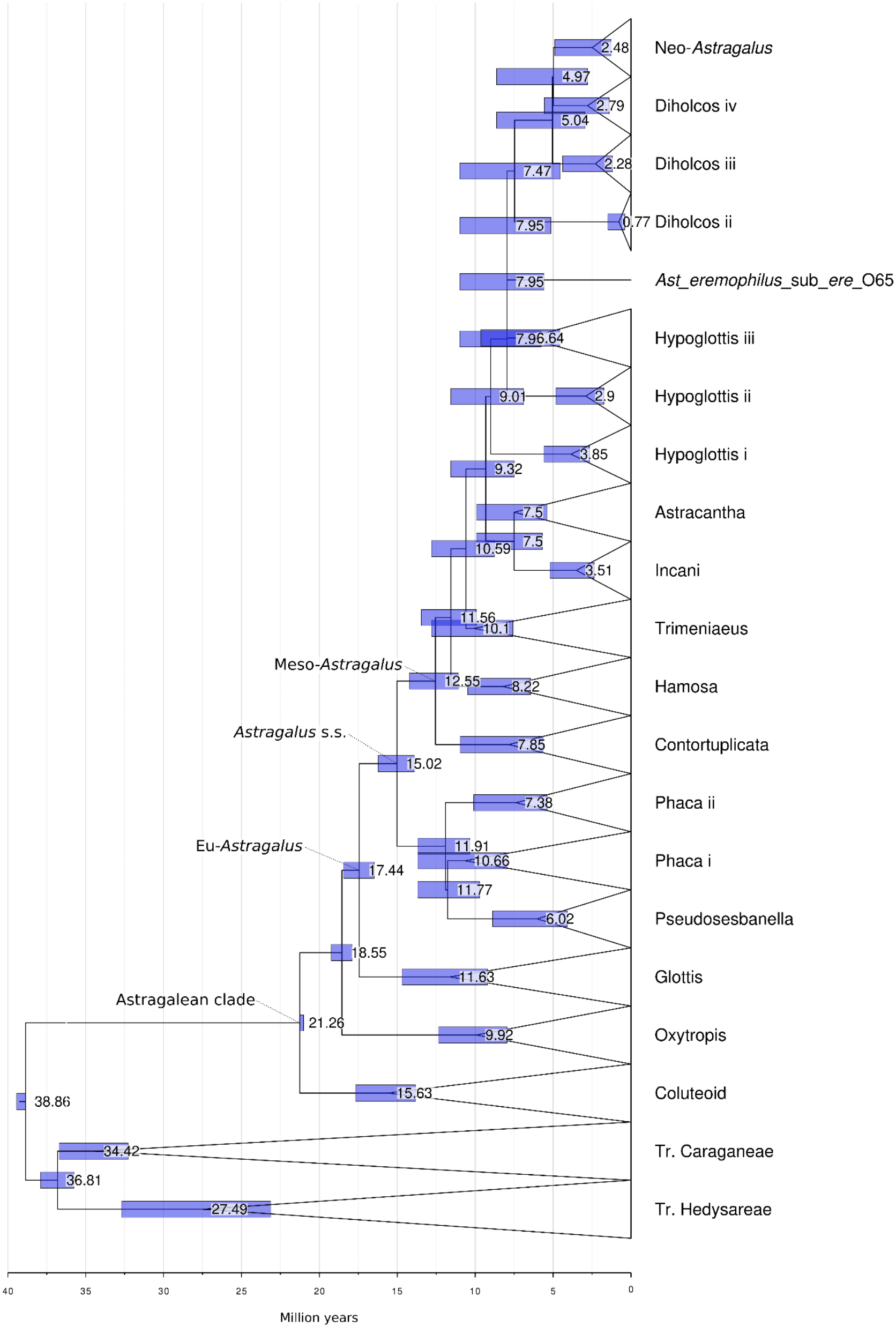
Dated phylogeny obtained using a large combined dataset which includes 370 samples (347 *Astragalus* species). Divergence times were calibrated using secondary age constraints at the root (38.84–40.21 Ma) and at the split between the Coluteoid clade and the remaining Astragalean clade (20.63–21.19 Ma). Blue bars represent 95% confidence intervals for node-age estimates; values indicate median divergence times in millions of years (Ma).

**Fig. S3.**
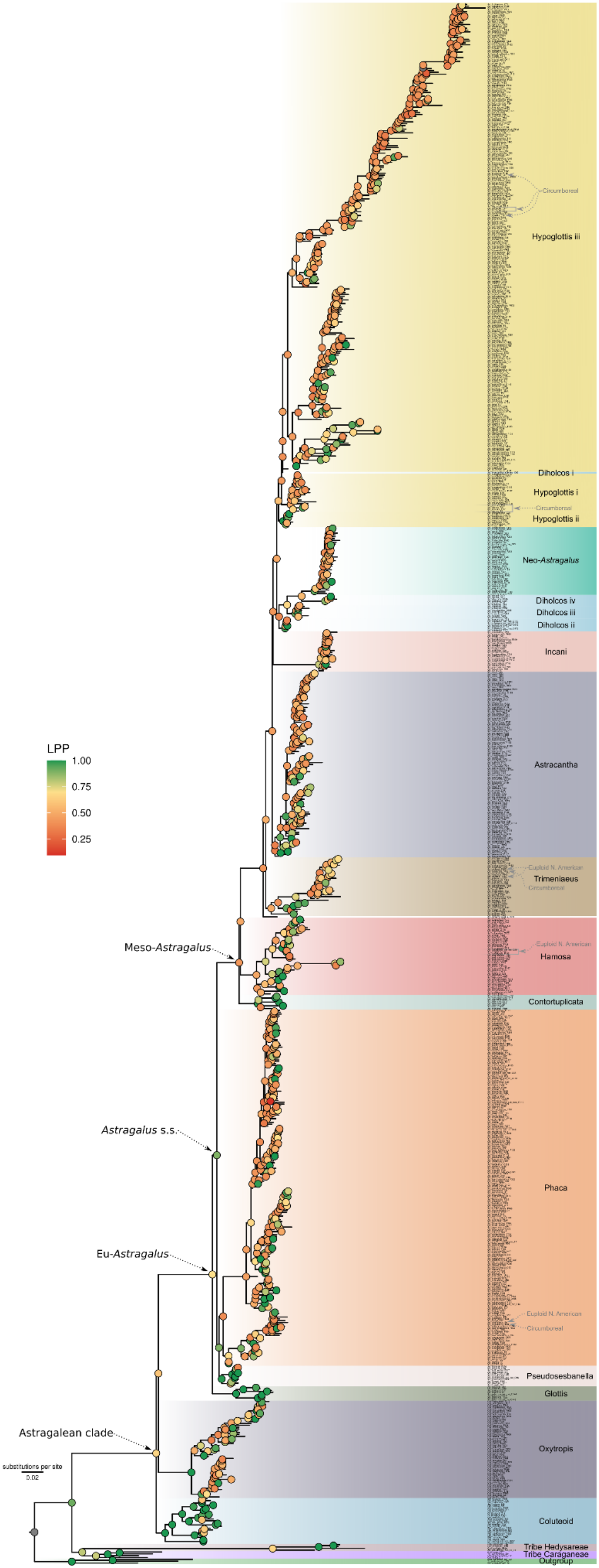
Coalescent-based phylogeny constructed with 831 homologous sequences belonging to 850 taxa, of which 760 *Astragalus* corresponds to 621 species.

**Fig. S4.**
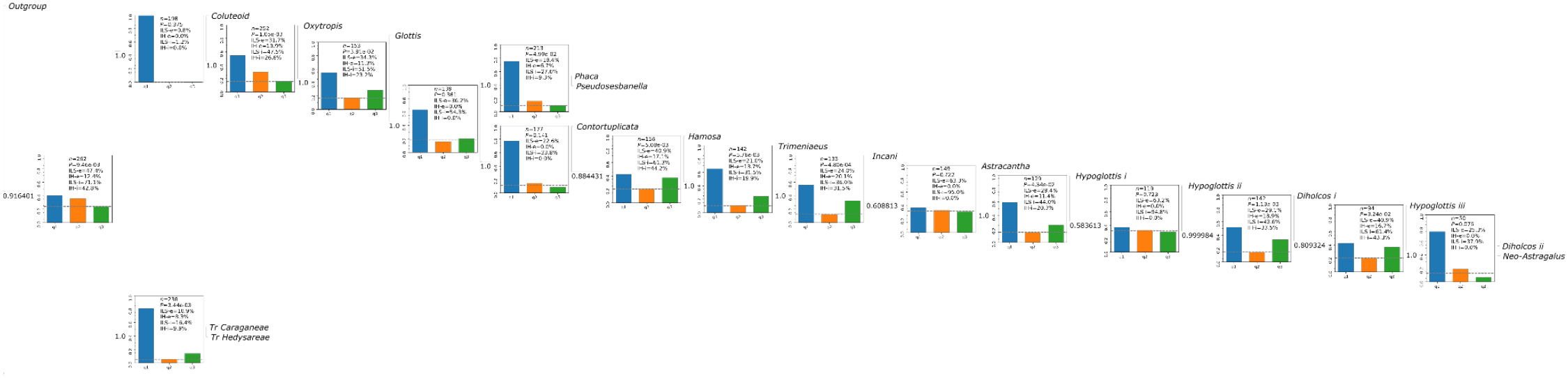
PhyTop results based on the coalescent species tree and collapsed by major clades along the backbone.

**Fig. S5.**
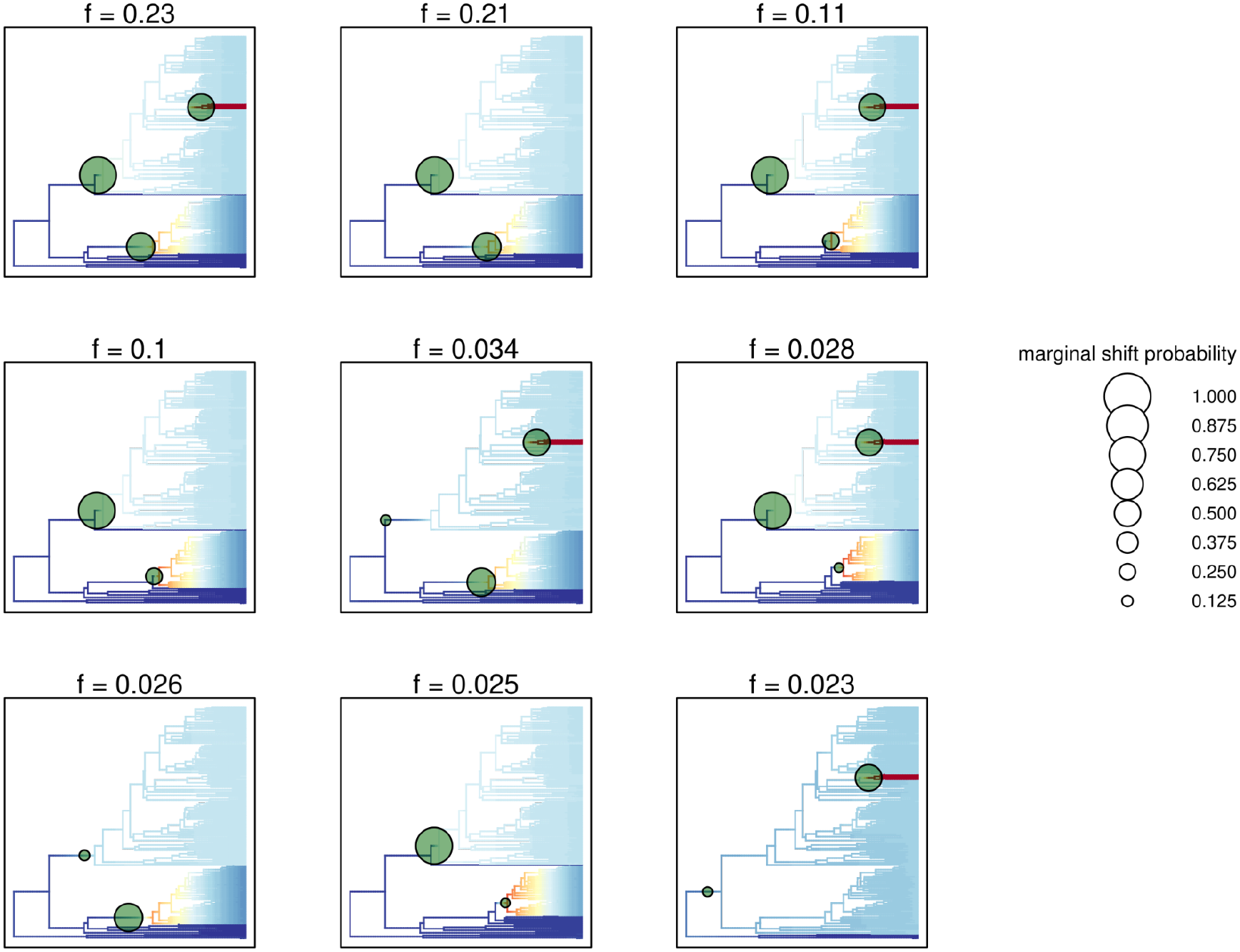
Credible shift set produced by BAMM analysis.

**Fig. S6.**
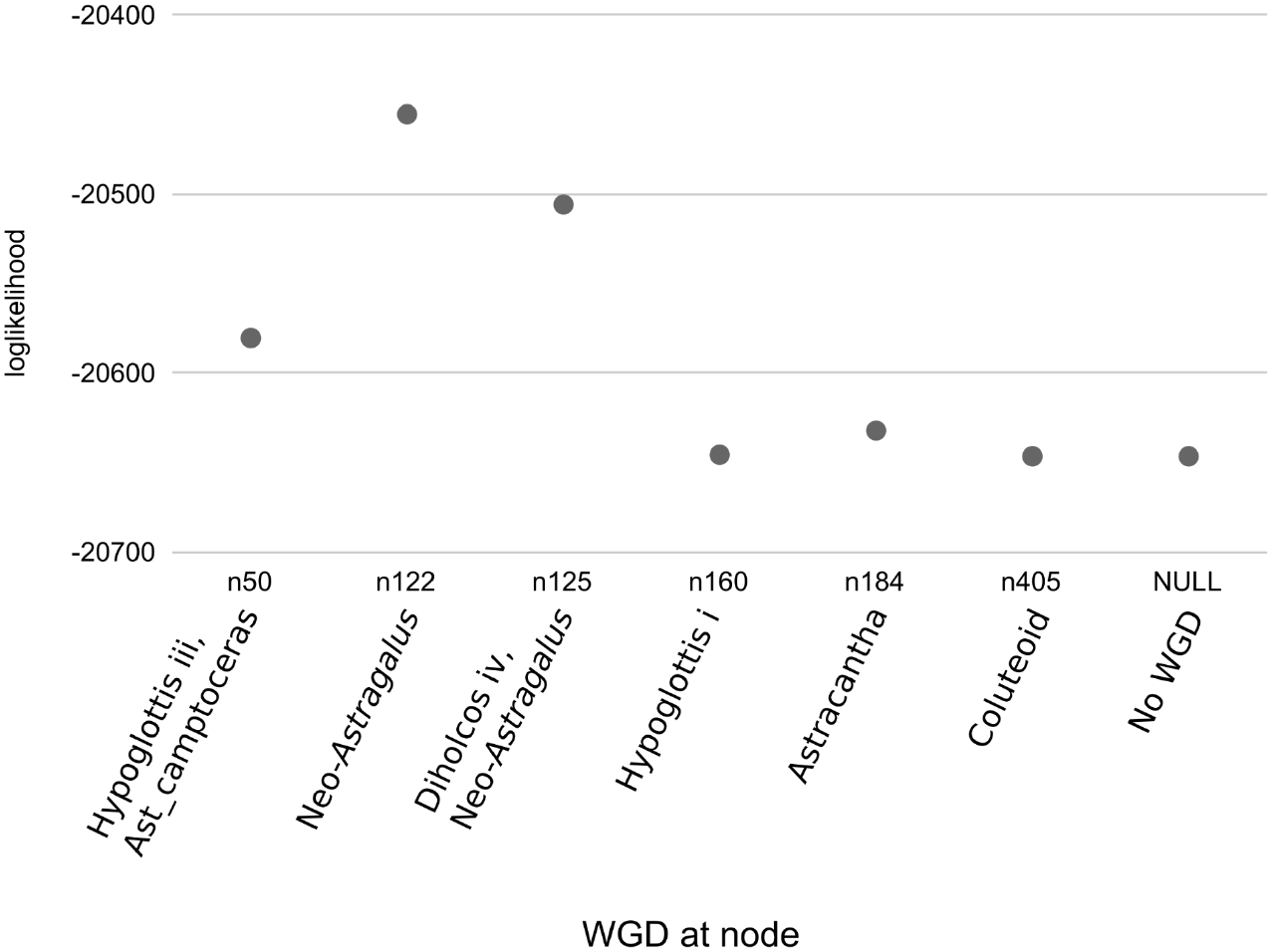
Likelihood of different WGD events occurring at a particular node inferred from gene counts using WGDgc.

## Supplementary tables

See supplementary table in excel format.

